# The brain simulates actions and their consequences during REM sleep

**DOI:** 10.1101/2024.08.13.607810

**Authors:** Yuta Senzai, Massimo Scanziani

## Abstract

Vivid dreams mostly occur during a phase of sleep called REM^1–5^. During REM sleep, the brain’s internal representation of direction keeps shifting like that of an awake animal moving through its environment^6–8^. What causes these shifts, given the immobility of the sleeping animal? Here we show that the superior colliculus of the mouse, a motor command center involved in orienting movements^9–15^, issues motor commands during REM sleep, e.g. turn left, that are similar to those issued in the awake behaving animal. Strikingly, these motor commands, despite not being executed, shift the internal representation of direction as if the animal had turned. Thus, during REM sleep, the brain simulates actions by issuing motor commands that, while not executed, have consequences as if they had been. This study suggests that the sleeping brain, while disengaged from the external world, uses its internal model of the world to simulate interactions with it.

---

While asleep, our brain generates virtual worlds we call dreams. Despite being largely cut off from the external world, the brain simulates interactions with it. In a dream we can act, and those actions have consequences as if they were executed in the physical world^2,16,17^. But what is an action without a movement and how can it have consequences if it is not executed? We explored the possibility that, while dreaming, the brain represents the consequences of issued motor commands, even if these motor commands are not executed. That is, that the brain simulates interactions with the world by using its internal model of the world. We decided to address this question by recording from the mouse brain during REM sleep, a phase of sleep which in humans is associated with perceptually vivid dreams^1^, and during which the execution of actions is prevented by muscle atonia^18^.

We chose the internal representation of heading as an experimental model. This system is made of neurons, so called head direction cells, whose ensemble activity reports the direction of the head of an animal along the azimuth, like an inner compass^19–21^. In awake animals, the action of turning the head leads to the activation of the vestibular system, which, as a consequence, contribute to updating of the brains’ representation of heading to match the new head direction^19,22^. Interestingly, during REM sleep the internal representation of heading keeps shifting, despite the immobility of the animal’s head^6,7^. These shifts resemble those observed in awake behaving animals and are referred to as shifts in “virtual heading”^6,8^. What triggers the shifts in the internal representation of heading during REM sleep? Do these changes in virtual heading simply represent spontaneous drifts of the head direction system or are they triggered by some other part of the brain? We tested for the potential contribution of the superior colliculus (SC), a midbrain structure involved in orienting movements^9,10^. During wakefulness, motor commands issued by the rodent’s SC trigger head turns^11–13,23,24^ and the resulting sensory consequences contribute to updating the internal representation of heading^19,22^. We reasoned that if the SC issues motor commands also during REM sleep, these commands may shift the internal representation of heading even in the absence of head turns and of their sensory consequences. In other words, we postulated the existence of an internal model that maps motor commands issued by the SC into shifts of the head direction system. To address this possibility, we first compared SC activity occurring during wakefulness with that occurring during REM sleep. We then tested whether SC activity during REM sleep predicts shifts in the virtual representation of heading, and finally perturbed the SC to establish causality between its activity and virtual heading.

## Turn-like activity in the superior colliculus during REM sleep

We performed extracellular recordings with linear probes from the intermediate and deep layers of the left SC of mice that were free to explore an open field arena while their head direction was monitored by a camera placed above the arena (Fig. 1a,b). Head turns were defined as changes in head direction whose speed was larger than 100 deg/s (see methods). We aligned the activity of SC neurons to the onset of clockwise (CW) or counterclockwise (CCW) head turns to obtain a peri-turn time histogram (PTTH; Fig. 1c,d). The activity of about half of the neurons was significantly modulated around the time of head turns in at least one direction (170 out of 350; n = 8 mice) and we refer to these neurons as turn cells. We quantified the turn direction preference of turn cells using the turn index (TI; the activity during CW head turns minus the activity during CCW head turns normalized by basal activity; Extended Data Fig. 1). The vast majority of turn cells had a positive TI (111 out of 170, Extended Data Fig. 1) i.e. they were CW turn cells, consistent with the recordings being performed in the left SC^13^ (Fig. 1d). To capture the temporal relationship in the activity of turn cells relative to each other we performed pairwise correlation analysis (Extended Data Fig. 2a,b). CW turn cells had a significantly larger probability of firing together as compared to CW-CCW pairs or CCW-CCW pairs (Extended Data Fig. 2c). To determine whether periods during which CW turn cells fire together correspond to an increased probability of the animal to perform CW head turns, we identified “population turn events”, that is, time intervals during which the combined activity of CW turn cells exceeded one standard deviation above the mean (Fig.1e). Indeed, during these population turn events, the probability of CW head turns increased 2.4 times (Fig. 1f). For each CW turn cell, we computed the spike-triggered average (STA) of head direction. This STA allows us to extract the average head turn amplitude around the time of a spike, i.e. the difference in head direction 200 ms before and after the spike (positive values represent CW turns; Fig. 1g). The average head turn amplitude around the time of a spike was significantly larger for spikes belonging to population turn events as compared to spikes outside of such events (Fig. 1g,h), consistent with increased probability of the animal to perform CW head turns during population turn events. Finally, the average head turn amplitude around the time of a spike was positively correlated with the TI (Extended Data Fig. 1). Thus, consistent with the role of SC as a motor command center for orienting movements, in freely moving animals, the activity of SC neurons predicts head turns, neurons with the same head turn direction preference are likely to fire together, and when several head turn neurons fire together, the likelihood of head turns is increased.

**Figure 1.**
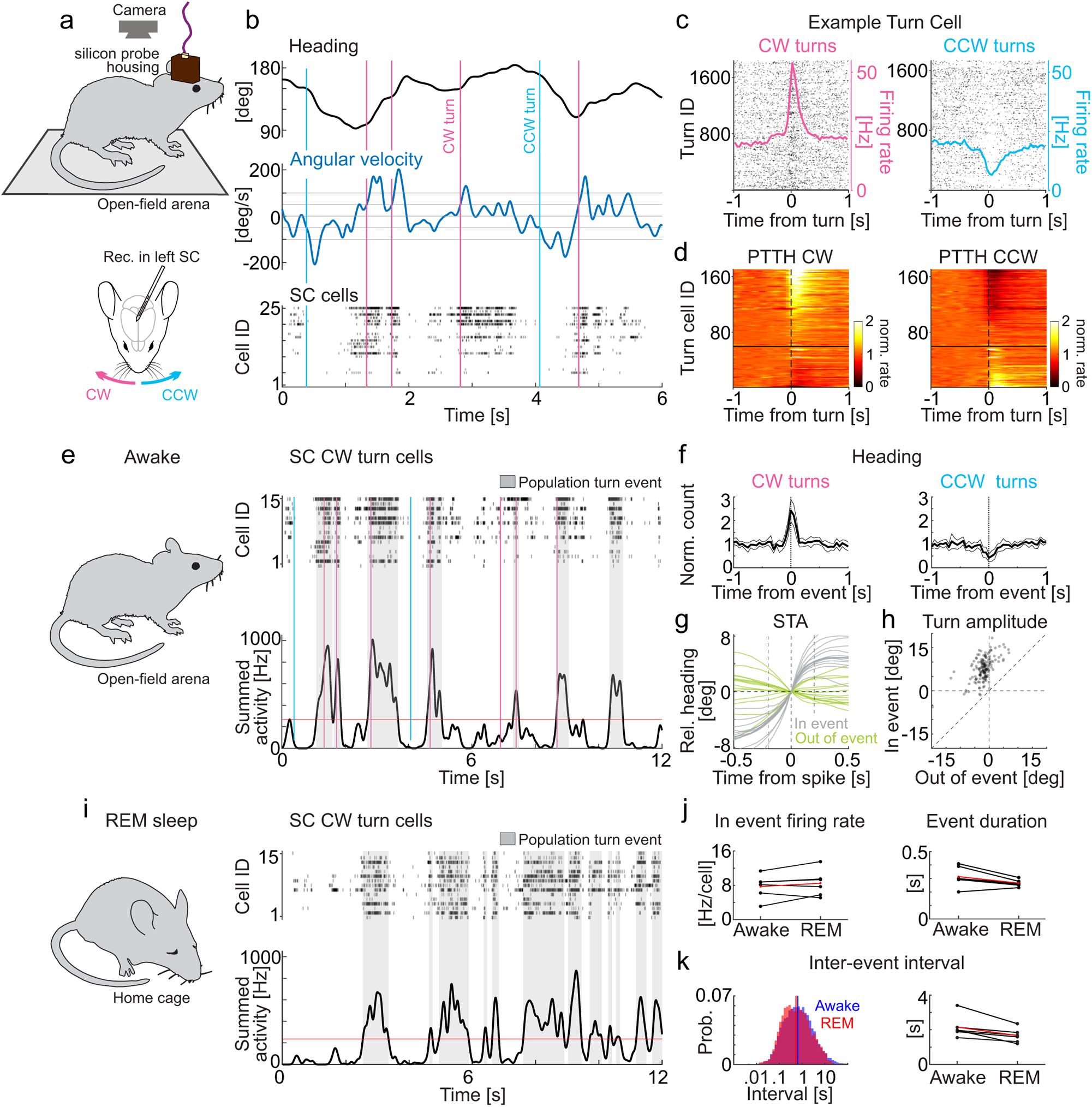
Turn-like activity in the superior colliculus during REM sleep. (a) Schematic illustration of the experimental configuration. The mouse, freely moving in an open-field arena is monitored with a top-view camera (top) while the electrical activity of the left superior colliculus (SC) is recorded with a chronically implanted linear probe (bottom). CW: clockwise head turn; CCW: counterclockwise head turn. (b) Top: Head direction (heading) of the animal along the azimuth over a 6 s period. Middle: Angular velocity of the head of the animal along the azimuth (time derivative of the top trace). Positive and negative values correspond to CW and CCW turns, respectively. The pink and blue vertical lines are the onset of CW and CCW turns, respectively. The horizontal lines at +/−100 deg/s are the thresholds for considering a head movement as a head turn. The horizontal lines at +/− 50 deg/s are the thresholds delimiting the onset and offset of the head turn. Bottom: Raster plot of the firing of the 25 SC neurons recorded during this example session. (c) Raster plot and superimposed peri turn time histogram (PTTH; continuous traces) of an example SC turn cell (#23 in b) aligned to the onset of approximately 1800 CW (left) and CCW (right) turns. (d) Normalized PTTHs of 170 turn cells for CW (left) and CCW (right) turns from 8 animals. The cells are ranked according to their turn index (see methods) with higher identity (ID) numbers corresponding to larger turn indices. The black horizontal line indicates the boundary between positive and negative turn indices, i.e. CW and CCW preferring cells, respectively. (e) Raster plot of 15 CW turn cells (top) and their summed activity (bottom) over a 12 s period. The first 6 s are the same as in (b). Vertical lines as in (b). The red horizontal line is the threshold above which the summed activity is considered a “population turn event”. The gray shaded area represents the duration of population turn events. Note that most CW head turns occur around the onset of a population turn event. (f) Normalized frequency of CW (left panel) and CCW (right panel) head turns at the onset of CW population turn events for 6 mice. Thick line is average; thin line is SD. (g) Spike triggered average (STA) of head direction for each of the 15 CW turn cells in (e). Gray traces are STAs for spikes inside a population turn event (In event). Light green traces are STAs for spikes outside population turn events (Out of event). To compute the STA the heading is set at 0 deg at the time of each spike such as to report the relative change in head direction. (h) Scatter plot of turn amplitudes, computed as the difference in the STA of head direction 200 ms before and after the spike (vertical dotted lines in (g)), for spikes in and out of population turn events for 156 cells in 6 animals (p = 2.4*10^−27^; signed rank test). (i) As in E but during REM sleep. Same 15 SC CW turn cells. Note the presence of population turn events. (j) The mean firing rate of cells during a population turn event was similar between wakefulness and REM sleep (left; p = 0.31; signed-rank test) as was the event duration (right; p = 0.05; signed-rank test). Black: 6 individual mice; Red: Population mean. (k) : The distribution of inter-event intervals was shifted towards shorter intervals during REM sleep (right; p = 2.8*10^−11^, KS test. left; blue awake; red: REM sleep) as was the mean inter-event interval (right; p= 0.03; signed-rank test). Black: 6 individual mice; Red: Population mean.

How does SC activity during wakefulness compare with that occurring during REM sleep? Animals were allowed to fall asleep in their home cage and REM sleep periods were identified using standard electrophysiological parameters (methods). We discovered that the activity of CW turn cells recorded in the sleeping animal was very similar to that observed in the awake animal. Not only were population turn events also present during REM sleep (Fig. 1i), but they occurred at a slightly higher frequency (Fig. 1k). Furthermore, the average duration of these events and the overall firing rate within those events were not significantly different from wakefulness (Fig. 1j). Crucially, the correlational structure of turn cells during REM sleep was similar to what was observed in awake animals (Extended Data Fig. 2c-g), with CW turn cells having a significantly larger probability of firing together as compared to CW-CCW pairs or CCW-CCW pairs (Extended Data Fig. 2f). Thus, during REM sleep, the activity of turn cells in SC resembles that observed in awake behaving animals performing head turns.

## The SC predicts virtual head turns during REM sleep

Does the activity in SC during REM sleep predict head turns? The head of the animal does not move during REM sleep, yet we can detect changes in virtual heading by recording from the head direction system^6^. We thus inserted a second linear probe in the anterodorsal nucleus of the thalamus (ADN; Fig. 2a) to record from head direction cells^25^ (Fig, 2b) whose population activity represents, during wakefulness, the actual heading of the animal^25^ and, during REM sleep, its virtual heading^6,8^. We used the actual heading of the awake animal, monitored with the camera above the arena, to train a decoder to report the heading based on ADN activity (Fig. 2c). The decoder, when tested on untrained periods, had an average error of 9.8 ± 1.3 degrees. We used the same decoder to report the virtual heading of the animal during REM sleep (Fig. 2c). The virtual heading shifts with dynamics that resembled those occurring during wakefulness, despite the immobility of the head^6,8^ and we refer to these shifts in virtual heading as “virtual head turns”. With this recording configuration we can align SC activity to the onset of head turns in the awake animal and to the onset of virtual head turns in the sleeping animal and compare the two.

**Figure 2.**
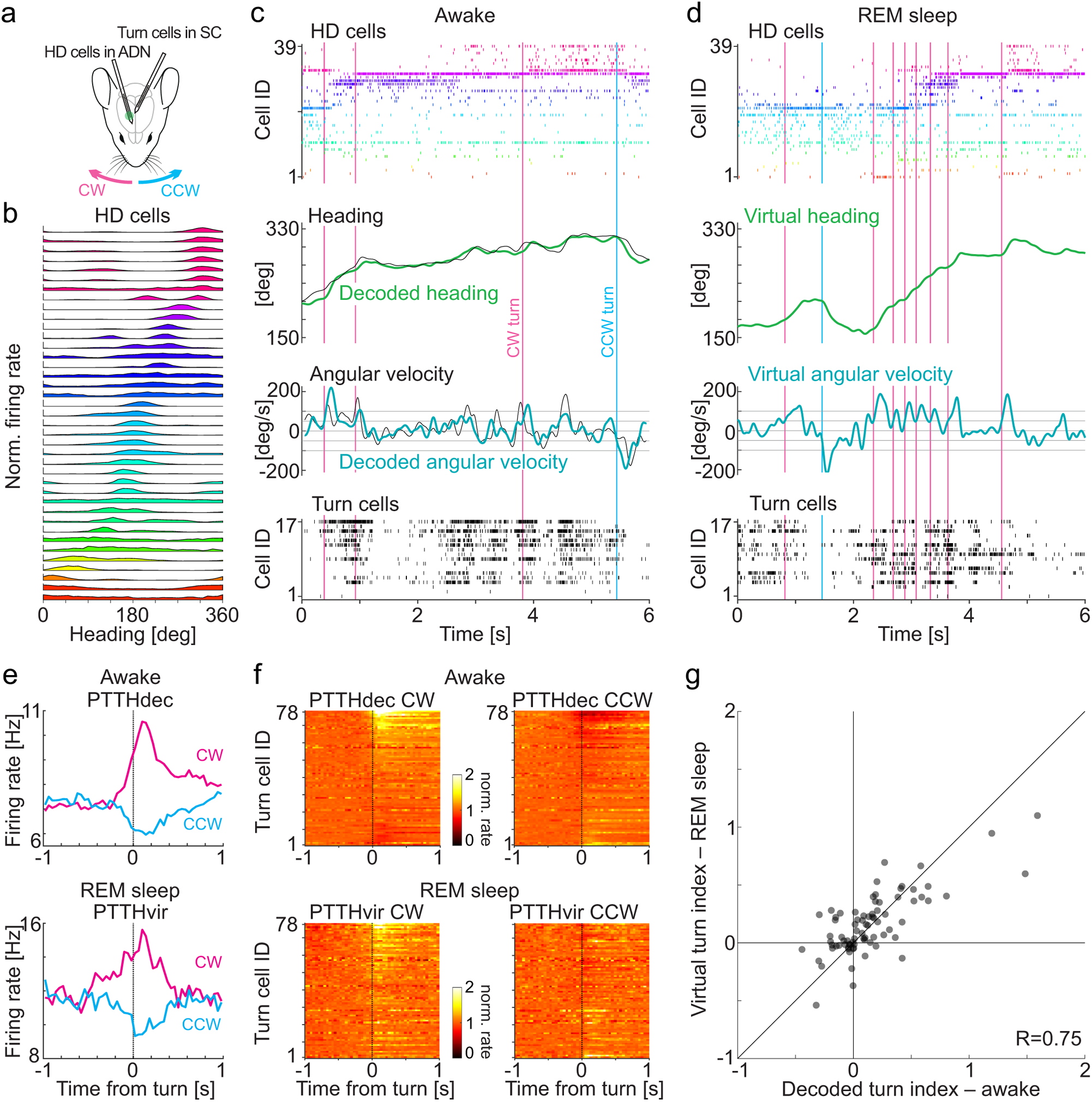
Turn-like activity in the superior colliculus during virtual head turns in REM sleep. (a) Schematic of the recording configuration from the left SC, to record turn cells, and the right anterodorsal thalamic nucleus (ADN) to record head direction (HD) cells. (b) Normalized tuning curves of 39 HD cells recorded in the ADN of an example recording session in an awake animal. Cells are ranked according to the similarity of their head direction preference. (c) Awake animal. Top: Raster plot of the 39 HD cells recorded in the ADN illustrated in (b) (same color code) during a 6 s example period. Second panel from top: Actual head direction (black) and decoded head direction (green; decoded from the 39 ADN cells shown in (b)). Third panel from top: Actual head velocity (black; time derivative of the actual head direction) and decoded head velocity (green; time derivative of decoded head direction). Horizontal lines as in Fig. 1b. The pink and blue vertical lines are the onset of decoded CW and CCW turns, respectively. Note that while the decoded head velocity misses a few actual head turns the detected ones occurred close to the actual ones. Bottom: raster plot of 17 turn cells recorded in SC (cell 1 and 2 are CCW, 3-17 are CW). (d) Same recording session as in (c) but during REM sleep. During REM sleep, the head direction and angular velocity decoded from the activity of HD cells are referred to as “virtual”. (e) PTTH of an example SC cell (#9 in (c)) aligned to the onset of decoded turns (PTTHdec) during wakefulness (top) and virtual turns (PTTHvir) during REM sleep (bottom). CW and CCW in pink and blue, respectively. Note the increase in firing rate of the cell around the time of a CW turn, irrespective of whether it is a decoded head turn during wakefulness or a virtual head turn during REM sleep. (f) Normalized PTTHs of 78 cells for CW (left) and CCW (right) turns from 5 animals. Top: PTTHdec during wakefulness. Bottom PTTHvir of the same cells during REM sleep. The cells are ranked according to their decoded turn index computed from their activity during wakefulness in the home cage (see also Suppl. Fig 3b). (g). Scatter plot of the turn index (TI) from the same 78 cells as in (f) computed in awake (decoded TI) against the TI computed during REM sleep (virtual TI). Note the significant correlation (P=2.4*10^−15^).

We computed the PTTH of individual turn cells obtained during wakefulness and REM sleep. The PTTH obtained during REM sleep is necessarily computed by aligning the activity of turn cells to the onset of virtual head turns and we refer to it as PTTHvir. For comparative purposes, the PTTH obtained during wakefulness was computed by aligning the activity of turn cells to the onset of decoded head turns rather than actual head turns and we refer to it as PTTHdec, to distinguish it from the PTTH obtained from actual head turns (Extended Data Fig. 3a). Strikingly, the activity of a large fraction of turn cells in SC was significantly modulated around the time of virtual head turns (61 out of 79; n = 5 mice) and, like turn cells recorded in the awake animal, the majority of those cells preferred CW than CCW virtual head turns (Fig. 2e,f).

Do SC turn cells keep the same turn direction preference between wakefulness and REM sleep? We compared the TI obtained during wakefulness with that obtained during REM sleep. The TI obtained during REM sleep (TIvir) was computed as the difference in the activity of turn cells during CW and CCW virtual head turns, normalized by basal activity. Again, for comparative purposes, the TI obtained during wakefulness was computed from decoded rather than actual head turns, and we refer to it as TIdec (consistent with our decoding accuracy, TI and TIdec were very similar (Extended Data Fig. 3b)). Crucially, there was a strong correlation between TIdec and TIvir (Fig. 2g), indicating that SC turn cells maintain similar turn preferences for actual head turns performed by awake behaving animals and virtual head turns performed during REM sleep.

How similar are actual and virtual head turns predicted by the activity of SC turn cells? For each turn cell we computed the STA of head direction during wakefulness and during REM sleep. We used the STA of virtual head direction (STAvir) obtained during REM and the STA of decoded head direction (STAdec) obtained during wakefulness to extract the average amplitude of a head turn around the time of a spike (Fig. 3a). The average amplitude of virtual and decoded head turns around the time of a spike in an SC turn cells were significantly correlated (Fig. 3b). Furthermore, during REM sleep, the average head turn was larger for spikes belonging to a population turn event as compared to those outside population turn events, as observed in awake animals (Fig. 3c,d). Taken together, these data show that the activity of turn cells in SC predicts head turns represented by the head direction system, irrespective of whether the animal is awake or asleep.

**Figure 3.**
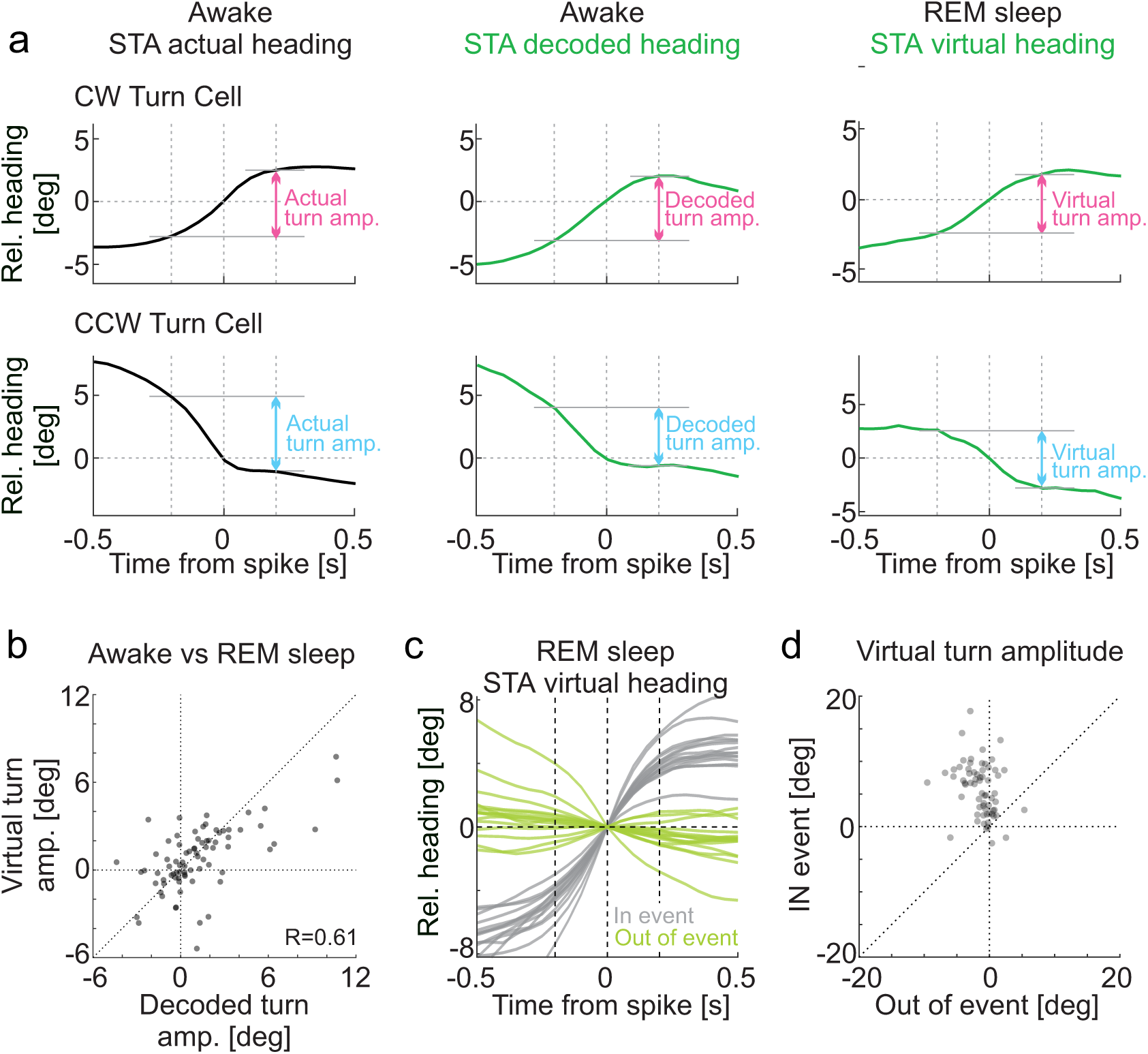
Activity in the superior colliculus predicts virtual head turns during REM sleep. (a) Top row: Example CW turn cell; Bottom row: Example CCW turn cell. Left: Spike triggered average (STA) of actual head direction. Middle: STA of decoded head direction during wakefulness. Right: STA of virtual head direction during REM sleep. To compute the STAs the heading is set at 0 deg at the time of each spike such as to report the relative change in head direction. The turn amplitude (actual, decoded and virtual; double headed arrows) is the difference in head direction 200 ms before and after the spike (vertical dotted lines). (b) Scatter plot of decoded turn amplitudes during wakefulness against virtual turn amplitudes during REM sleep for 78 turn cells (5 animals). Note the significant correlation (p=2.8*10^−9^). (c) STA of virtual head direction during REM sleep for the same 15 CW turn cells shown in Fig.1g. Gray traces are STA for spikes inside a population turn event. Light green traces are STA for spikes outside of a population turn event. (d) Scatter plot of virtual turn amplitudes computed for spikes in and outside of population turn events for 65 cells in 3 animals. Note that, like in the awake animal (Fig. 1h), during REM sleep, average turn amplitudes around spikes inside population turn events are larger than those outside of population turn events (p=2.6*10^−11^, signed rank test).

## The SC generates virtual head turns during REM sleep

Is there a causal relationship between the activity in SC and the representation of head direction in the ADN during REM sleep? In other words, is SC triggering virtual head turns in the ADN? We took advantage of the lateralization of orienting movements evoked by the SC, namely the fact that each SC is predominantly responsible for contraversive turns^12,13,23,24^ (i.e. left SC contributes to CW turns). We reasoned that if the SC contributes to the generation of virtual head turns recorded in the ADN during REM sleep, blocking the activity of the right SC while leaving left SC intact may lead to an increase in the number of CW relative to CCW virtual head turns. We thus recorded from head direction cells in the ADN and blocked activity in the right SC by injecting this structure with the sodium channel blocker tetrodotoxin (TTX; methods) while the animal was briefly head fixed for the duration of the procedure (Fig. 4a). Once the animal was placed back in its home cage, we waited for it to fall asleep. As soon as the animal entered REM sleep, the number of CW relative to CCW virtual head turns showed a significant increase compared to control conditions (Fig. 4b-d) leading to a concatenation of 360° CW virtual turns. A bias in the number of CW turns in untethered awake animals monitored after the recordings session, confirmed the action of TTX in the right SC (Extended Data Fig. 4). These data indicate a causal contribution of the SC in the generation of virtual head turns during REM sleep.

**Figure 4.**
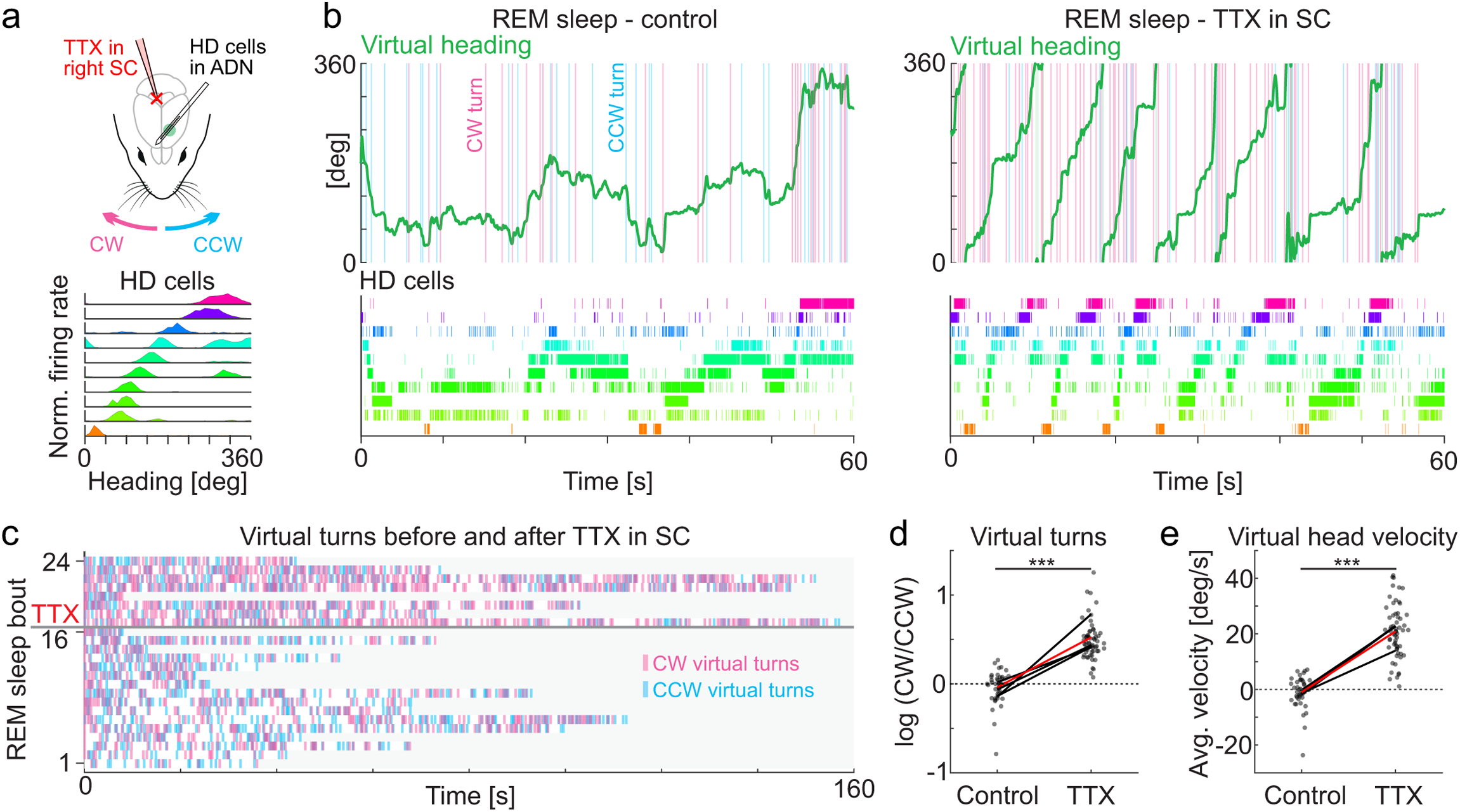
Activity in the SC generates virtual head turns during REM sleep. (a) Top: Schematic of the experimental configuration with a recording electrode in the left ADN to record head direction (HD) cells and a pipette to apply TTX to block activity in the right SC. Bottom: Normalized tuning curves of 10 example head direction cells of an example recording session in an awake animal. Cells are ranked according to the similarity of their head direction preference. (b) Top: Virtual heading (green trace) decoded during REM sleep under control conditions (left) and after TTX injection in the right SC (right). The pink and blue vertical lines are the onset of virtual CW and CCW turns, respectively. Bottom: Raster plots of the 10 HD cells illustrated in (a) used to decode the virtual heading (same color code). Note the strong increase in virtual CW turns with TTX in the right SC. (c) Raster plot of virtual CW (pink) and CCW (blue) turns during individual REM sleep bouts under control conditions (bout 1 to 16) and after injection of TTX in the right SC (bout 17 to 24). Same animal as in (b). (d) Population average of the logarithm of the ratio between the number of virtual CW and CCW turns during REM sleep in control versus TTX in SC. Black: 4 individual mice; Red: Population mean. Data points: individual REM sleep bouts (43 bouts in control and 62 bouts in TTX). Note the increase in the fraction of virtual CW turns in TTX (p=1.5*10^−17^; Kruskal-Wallis test). (f) Population data of virtual head velocity averaged over individual REM sleep bouts in control versus TTX. Black: 4 mice; Red: Population mean. Data points: average virtual head velocity of individual REM sleep bouts (43 bouts in control and 62 bouts in TTX). Note the increase in positive velocities (i.e. CW turns) with TTX (p=9.1*10^−17^; Kruskal-Wallis test).

## Discussion

This study shows that during REM sleep motor commands issued in the SC trigger shifts in the representation of head direction in the ADN, despite the absence of head movements, and thus in the absence of sensory feedback from the vestibular canals. In other words, during REM sleep, the brain simulates an interaction with the external world by representing the consequences of a motor command.

How a motor command issued in SC leads to a virtual turn in the ADN during REM sleep remains to be established. The absence of head movements rules out an involvement of the vestibular organs. Furthermore, the absence of proprioceptive organs in the mouse external eye muscles^26^ also precludes the possibility that proprioceptive feedback from the eye movements that characterize REM sleep substantially contribute to shifts in the representation of heading. Thus, our data suggest that SC activity leads to virtual head turns by acting internally, possibly via efference copies, on those neuronal circuits that compute head direction. There are no monosynaptic projections linking the SC to the ADN. However, because motor commands from the SC are broadcast to the brainstem^24,27–30^ and the cortex^31,32^, and the ADN receives ascending inputs from the brain stem and descending inputs from the cortex^33^, the SC could use either of these routes to shift the representation of head direction.

Our data suggest that the simulations occurring during REM sleep are quite accurate. First, the activity of turn cells in SC during REM sleep is clustered in population turn events similar to what observed in awake behaving animals when SC activity triggers actual head turns (Fig. 1i-k). Second, the activity of SC turn cells is similarly modulated around the time of actual head turns in awake behaving animals and around the time of virtual head turns in sleeping animals (Fig. 2e-g). Third, like in wakefulness, during REM sleep, predicted shifts in head directions were larger for spikes belonging to population turn events (Fig. 3c,d). Thus, the putative internal model that links motor commands issued by the SC to a shift in the representation of head direction in the ADN during REM sleep reproduces the consequences of an actual head turn in the awake behaving animal. Whether the dynamics of the turn are also preserved during these simulations will be resolved with future studies.

Studies of dream content in humans support the idea that the simulations occurring during REM sleep result from the unfolding of the subject’s internal model of the world^16,17^. Furthermore, given that rapid eye movements during REM sleep likely reflect some of the cognitive processes occurring in the sleeping brain ^3,8,34,35^, the fact that human patients with unilateral neglect perform lateralized rapid eye movements during REM sleep^36,37^ is consistent with the possibility that the same internal models that shape the brain’s activity in wakefulness emerge during REM sleep.

Finally, it has been postulated that REM sleep itself contributes to the formation and refinement of internal models^16,38^. It is thus conceivable that the internal model that coordinates activity between SC and ADN observed during REM sleep may be at work also in the awake animal. This would allow the brain to predict shifts in head direction following the issuing of a motor command in the SC, before the motor command is executed and thus have a real time estimate of heading rather than depending on the necessarily delayed sensory feedback from the vestibular canals. If so, however, the difference between the predicted shift in head direction and the actual shift would have to be signaled to correct the representation of head direction from possible errors in prediction. It is interesting to consider that, during REM sleep, such error signals must be muted^17^, otherwise deviations of virtual head direction from actual head direction would be continuously corrected. In the absence of error signals, these internal models are free to unfold and may represent the building blocks of the brain’s generative activity during REM or REM-like sleep in humans and animals^39–41^.

## Acknowledgments

We thank Pooja Saraf, Qui Ying Wu and Leo Ruan for technical assistance, Loren Frank and Eric Denovellis for sharing their decoding method, Brenda Bloodgood for critical reading of the manuscript and the former and current members of the Scanziani lab for discussions.

## Author contributions

Y.S. and M.S. designed the study. Y.S. conducted all experiments and analyses. Y.S. and M.S. wrote the paper.

## Materials and Methods

### Animals

All experimental procedures were conducted in accordance with the regulations of the Institutional Animal Care and Use Committee (IACUC, AN179056) of the University of California, San Francisco. All mice were housed on a reversed cycle (light/dark cycle 12/12 h) with free access to food. Data were collected from male or female C57BL/6J mice (Jackson Laboratory #000664). At the start of the experiments, all mice were between 2.7 and 12 months old.

### Surgery and electrode implantation

Mice were implanted under 2% isoflurane anesthesia. The procedure was performed in two phases on two distinct days: First, the implantation of the headplate, and second, the implantation of the silicon probes together with, when applicable, the craniotomy for TTX injections. The headplate base was placed on the skull with dental cement (Unifast LC, GC America; OptiBond Universal, Kerr Dental). For some animals, the headplate base was integrated with infrared LEDs and 3D-printed holders for eye-tracking cameras^8^. After a minimum post-op recovery of five days in their home cage, mice were habituated to freely explore the open field arena (see below) with cables for electrophysiology attached to their head, but not electrically connected, in order to become familiar with the recording apparatus. Following habituation to the experimental setup, mice underwent the second surgical phase, namely the implantation of the silicon probes. A silicon probe was mounted on a movable microdrive to record from the anterodorsal nucleus of the thalamus (ADN) or the superior colliculus (SC). The types of silicon probes used included: Single-shank probes (Diagnostic Biochips P64-1, 64 channels per shank, 20 μm interchannel pitch); two-shank probes (Diagnostic Biochips P64-3, 250 μm inter-shank distance, 32 channels per shank, 25 μm interchannel pitch), and four-shank probes (Diagnostic Biochips P64-4, 250 μm inter-shank distance, 16 channels per shank, 20 μm interchannel pitch). Each of the shanks of the probe were inserted, through a craniotomy, at the coordinates described in Table S1. The probes were initially lowered to 0.8-1.5 mm below the brain surface. After recovery from isoflurane anesthesia, the probes were lowered to the ADN or the SC by moving the microdrive over one to three days. A ground electrode (0.005” diameter stainless wire, A-M systems) was inserted in the cerebellum. During the second phase of surgery, for mice undergoing TTX injections in the right SC, a craniotomy was made at the following coordinates relative to Lambda: anteroposterior: +0.2 mm, mediolateral: +1.4 mm. This craniotomy was sealed with biocompatible silicone sealant and removed just before the TTX injection (World Precision Instruments, Kwik-Cast). Experiments were performed after a minimum of 6 hours since the last probe depth adjustment.

### Extracellular electrophysiological recordings

Recording sessions from the ADN and/or the SC, occurred both while mice explored the open field arena for food as well as when they were in their home cage, and lasted for 60-120 min. The entire recording amounted to a total of 10 to 22 hours per mouse. Electrophysiological data were acquired using an Intan RHD2000 system (Intan Technologies LLC) band-pass filtered between 0.1 Hz and 7.5 kHz and digitized at 20 kHz. Spike sorting was performed semi-automatically, using Kilosort 2.0 ^42^. This was followed by manual adjustment of the waveform clusters using the software Phy, which was performed blinded to other data. For the local field potential (LFP), the wide-band signal was down-sampled to 1.25 kHz.

### Open field arena

The rectangular open field arena (50 × 28 cm) was surrounded by 25 cm high walls displaying salient visual cues. The arena base and the walls were made from white plastic. The arena was illuminated by visible room lights as well as by infrared LED lights for video acquisition. The video (50 Hz sampling rate) was captured by a CMOS camera (Basler acA1300-200um) placed above the arena with an infrared pass filter (Hoya) in front of the lens. The heading of the animal was detected based on the vector connecting the neck to the nose of the mouse. The extraction of the coordinates of the nose and the neck was performed by DeepLabCut ^43^. The heading was up-sampled to 1000 Hz by spline interpolation and smoothed with a Gaussian kernel with a standard deviation of 80 ms.

### Detecting head turns

We detected head turns by applying thresholds for angular head velocity. Angular head velocity was calculated from the outputs of DeepLabCut. Head turns were defined as head movements along the azimuthal plane whose angular velocity exceeded 100 deg/s. The beginning and the end of each head turn were defined as the first and the last time point, respectively, at which the angular head velocity exceeded 50 deg/s. To avoid spurious detection of head turns caused by glitches in DeepLabCut, we removed head turns with maximum head velocity over 500 deg/s from the further analysis.

### Turn Cells

We defined Turn Cells in the SC as neurons whose firing rate was significantly modulated by either CW or CCW head turns in the open field arena (Extended Data Fig. 1A). For this, we compared the average firing rate during the turn (i.e. the firing rate averaged over a 200 ms window starting at the onset of the turn) with the baseline firing rate (defined as the firing rate averaged over a period of 800 ms ending 200 ms before the onset of the turn). This analysis included at least 500 turns for each neuron. Significance (p < 10^−6^) was calculated using a paired t-test.

### Turn Index

We computed the Turn Index for each neuron from the peri-turn time histogram (PTTH) for CW or CCW head turn. The turn index was defined as the difference between the baseline-normalized firing rates of the neuron to CW and CCW turns (Extended Data Fig. 1a-c). The baseline-normalized firing rate was computed as the ratio of the average firing rate during the turn (i.e. the firing rate averaged over a 200 ms window starting at the onset of the turn) to the baseline firing rate (defined as the firing rate averaged over a period of 800 ms ending 200 ms before the onset of the turn). Turn Cells with a positive Turn Index were defined as CW Turn Cells, while Turn Cells with a negative Turn Index were defined as CCW Turn Cells.

### Detecting population turn events

Population turn events were defined as periods during which the summed activity of simultaneously recorded CW Turn Cells in the left SC exceeded the threshold of the mean plus one standard deviation. The mean and standard deviation were calculated separately for each brain state or recording condition, namely, wakefulness in the open field arena, wakefulness in the home cage, and REM sleep in the home cage. The beginning and the end of each population turn event, i.e. the duration, was defined as the first and the last time point, respectively, at which the summed firing rate of the CW Turn Cells exceeded the threshold. Inter-event intervals were calculated as the interval time from one event onset to the next event onset. For the spike-triggered average (STA) turn analysis, the spikes inside and outside the Turn Cell population events were separately analyzed. We defined the outside-event period as the period when the summed firing rate of CW Turn Cells is below the mean + 0.5 standard deviation.

### Brain state scoring

We identified wakefulness, non-REM sleep and REM sleep using the approach described in Watson (2016)^44^. Three metrics were extracted from the LFP data: power in the low-frequency band (<20 Hz), theta-ratio and electromyogram (EMG). To obtain the power in the low-frequency band, characteristically high in non-REM sleep, we first constructed spectrograms with a 1 s sliding 10 s-window Fast Fourier Transform (FFT) of the LFP at log-spaced frequencies between 1 and 100 Hz. We then performed principal components analysis (PCA) on the z-transformed (1-100 Hz) spectrogram and used the first PC as the power in the low-frequency band. The first PC mainly accounts for the variance in the spectrogram resulting from the alternation between “synchronous/low-frequency” (non-REM sleep) and “asynchronous/high-frequency” (Wakefulness/ REM sleep) states^44^. The theta-ratio was calculated as the ratio of 5-10 Hz power to 2-16 Hz power. EMG was extracted by detecting the zero time-lag correlation coefficients between 300-600 Hz filtered signals recorded at all sites^44^. Non-REM sleep periods were detected as intervals with high power in the low-frequency band. Among the remaining intervals, REM sleep periods were identified as those with weaker EMG values and stronger theta-ratio and the rest was wakefulness. Thresholds for each of the three metrics were set at the trough of the bimodal distribution of each metric^44^. After automated brain state scoring, all states were manually reviewed by the experimenter and minor corrections were made when discrepancies between automated scoring and user assessment occurred^44^.

### HD cells

The tuning of HD cells was computed as the ratio between histograms of spike count and the total time spent in each direction in bins of 1 degree. The tuning of each HD cell was tested for non-uniformity with a Rayleigh test. Neurons with significant non-uniformity (p<0.01) were identified as HD cells.

### Decoding of heading

Heading was decoded from the population activity of HD cells with a Bayesian decoding method that also integrates the information from both past and future spikes, using a package (replay_trajectory_classification) developed by Dr. Eric Denovellis^45^.

We set the decoding mode to “continuous movement” dynamics, which was modeled by a random walk transition matrix with a 2-degree standard deviation. Decoding was done using an “acausal algorithm” meaning that the decoded heading in the previous time bins and the following time bins influence the decoded heading at the current time bin through the random walk transition matrix. A 1-ms time bin and 1-degree heading bin were used to allow for high-resolution decoding. The data in the open field arena were split into two; the first half for training and the second half for testing. We used training data, i.e. firing rate of each HD cell in 1-ms time bins and the heading angle in 1-degree resolution in 1-ms time bins, to train the decoder. The decoder outputs the posterior probability of heading in 1-ms time bins. The heading with the highest posterior probability was chosen as the decoded heading. The decoder was applied to the testing data to evaluate the decoding accuracy.

### Detecting decoded and virtual head turns

We detected decoded head turns during wakefulness and virtual head turns during REM sleep by using the heading decoder outputs, i.e. decoded heading during wakefulness and virtual heading during REM sleep, respectively. First, we smoothed the decoded (or virtual) heading with a Gaussian kernel with a standard deviation of 80 ms. Then, the decoded (or virtual) head velocity was calculated by taking the first-order derivative of the smoothed heading. Like actual head turns (see “Detecting head turns”, above), decoded (or virtual) head turns were detected using a threshold of > 100 deg/s. The beginning and the end of each head turn were defined as the first and the last time point, respectively, at which the velocity of the decoded (or virtual) head velocity exceeded 50 deg/s.

### Decoded Turn Index and Virtual Turn Index

We computed the peri-turn time histogram (PTTH) for CW and CCW virtual head turns during REM sleep in the home cage, which we call PTTHvir. From PTTHvir, we computed the virtual Turn Index (TIvir) in the same way as computing TI from PTTH during the open field arena exploration (see “Turn index”, above). Similarly, PTTH for decoded head turns during wakefulness in the home cage (PTTHdec) and decoded Turn Index (TIdec) were computed.

### Pharmacology

Silencing of the right SC was performed by injecting 300 nl of 300 μM tetrodotoxin (TTX) at a speed of 30 nl min–1, using a beveled glass pipette (tip diameter, ∼20–40 μm) on a microinjector (Nanoject II, Drummond) with a Micro4 controller (World Precision Instruments), while mice were head-fixed. The injector was mounted on a micromanipulator (Luigs & Neumann) for stereotactic injection. Before the TTX injection, the Kwik-Cast sealant was removed from the craniotomy (anteroposterior: +0.2 mm, mediolateral: +1.4 mm) and the glass pipette was lowered, with a 30-degree angle toward medial, to 1350 μm depth below the brain surface. TTX was co-injected with the fluorescent dye Alexa594, for *post hoc* verification of the injection site. Following the 10 min long TTX injection and a 5 min post-injection wait time, mice were transferred to the open field arena or the home cage for recordings. Suppression of activity in the right SC was confirmed, for every experiment, by evaluating the turning behavior of untethered mice in the open field arena over a period of 10-15 min, 6-14 hours after the injection of TTX

### Histological processing

For anatomical analysis, mice were perfused transcardially with phosphate-buffered saline (PBS) and then with 4% paraformaldehyde (PFA) in PBS. Brains were extracted from the skulls, post-fixed in 4% PFA overnight at 4°C, and subsequently cut with a vibratome to 80-100 μm thick sequential coronal sections. Slices were collected and mounted in ProLong Gold (Life Technologies) or Vectashield mounting medium containing DAPI (Vector Laboratories H1500). Bright-field and fluorescence images were acquired using a Nikon Ti Inverted Microscope equipped with a Nikon DS-Qi2 monochrome camera (UCSF Nikon Imaging Center, 6D). The probe tracks were detected based on the gliosis formed around the probe.

### Quantification and statistical analysis

All statistical analyses were performed with standard MATLAB (MathWorks) functions. No specific analysis was used to estimate minimal population samples but the number of Animals and recorded cells were larger or similar to those employed in previous works^6,8^. Kruskal-Wallis one-way analysis of variance, Kolmogorov-Smirnov test, Rayleigh test, and paired t-test were used. Correlations were computed using Pearson’s correlation coefficient. Unless stated otherwise, all values in the text are given as mean average ± standard deviation.

### Inclusion criteria

For the decoding of virtual heading during REM sleep, we only included experiments that fulfilled the following two criteria: 1. At least 10 head direction cells were recorded. 2. The decoding of the actual head direction had an error of less than 20 degrees. For detecting population turn events, we included only mice from which we recorded more than 5 CW Turn Cells. For TTX injection experiments, we only included mice that, 6-14 hours following the TTX-injection, showed a biased turn behavior in the open field arena.

---

**Extended Data Figure 1.**
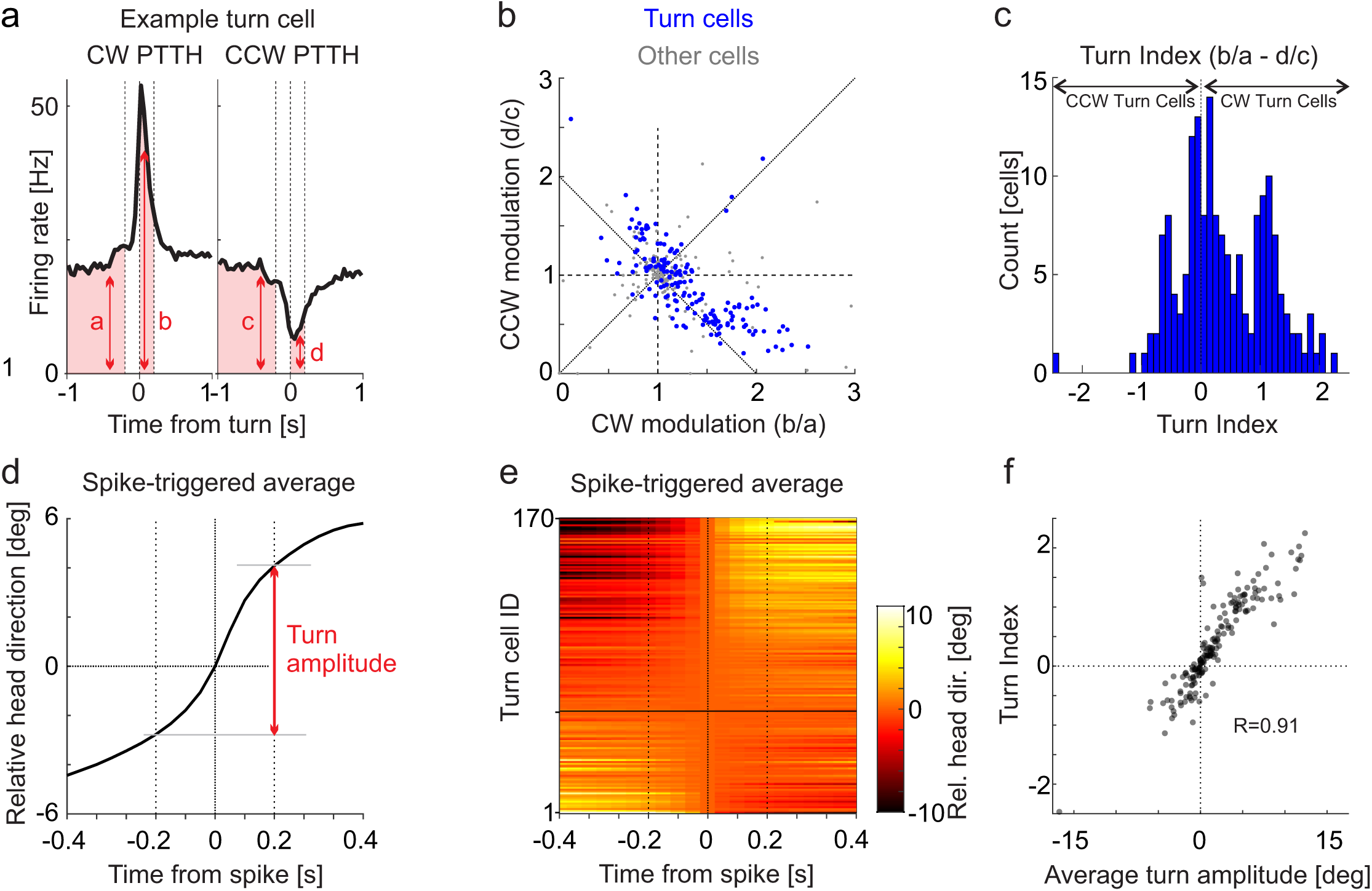
Turn Index and Spike Triggered Average of Head Direction. (a) Variables for calculating the Turn Index. Peri-turn time histogram (PTTH) of an example turn cell for clockwise (CW; left panel) and counterclockwise (CCW; right panel) turns (same cell as in Fig. 1c). Variables: a: baseline firing rate, averaged between 1 and 0.2 s before onset of CW turn. b: peak firing rate, averaged between 0 and 0.2 s after onset of CW turn. c: baseline firing rate, averaged between 1 and 0.2 s before onset of CCW turn. c: peak firing rate, averaged between 0 and 0.2 s after onset of CCW turn. Turn Index = (b/a) – (d/c) (b) Scatter plot of the modulation of the activity of SC neurons around the time of CW (b/a) against CCW (d/c) head turns. Turn cells. i.e. cells whose activity is significantly modulated around the time of head turns in at least one direction are in blue (n=170) and the rest in gray (n=180; 8 animals). (c) Distribution of Turn Indices for all turn cells (blue data points in (b)) Positive and negative turn indices define CW and CCW preferring turn cells, respectively. All cells were recorded in the left SC. (d) Calculating the average turn amplitude around the time of a spike in a turn cell. Spike triggered average (STA) of head direction from example turn cell. The heading is set at 0 deg at the time of each spike such as to report the relative change in head direction. The average turn amplitude (red vertical double headed arrow) is the difference in head direction 200 ms before and after the spike (vertical dotted lines). (e) STAs the 170 turn cells (8 animals). The cells are ranked according to their turn index. The black horizontal line indicates the boundary between positive and negative turn indices, i.e. CW and CCW preferring cells, respectively. (f) Average turn amplitude around the time of a spike and turn index are strongly correlated (P=1.1*10^−67^; n=170 cells).

**Extended Data Figure 2.**
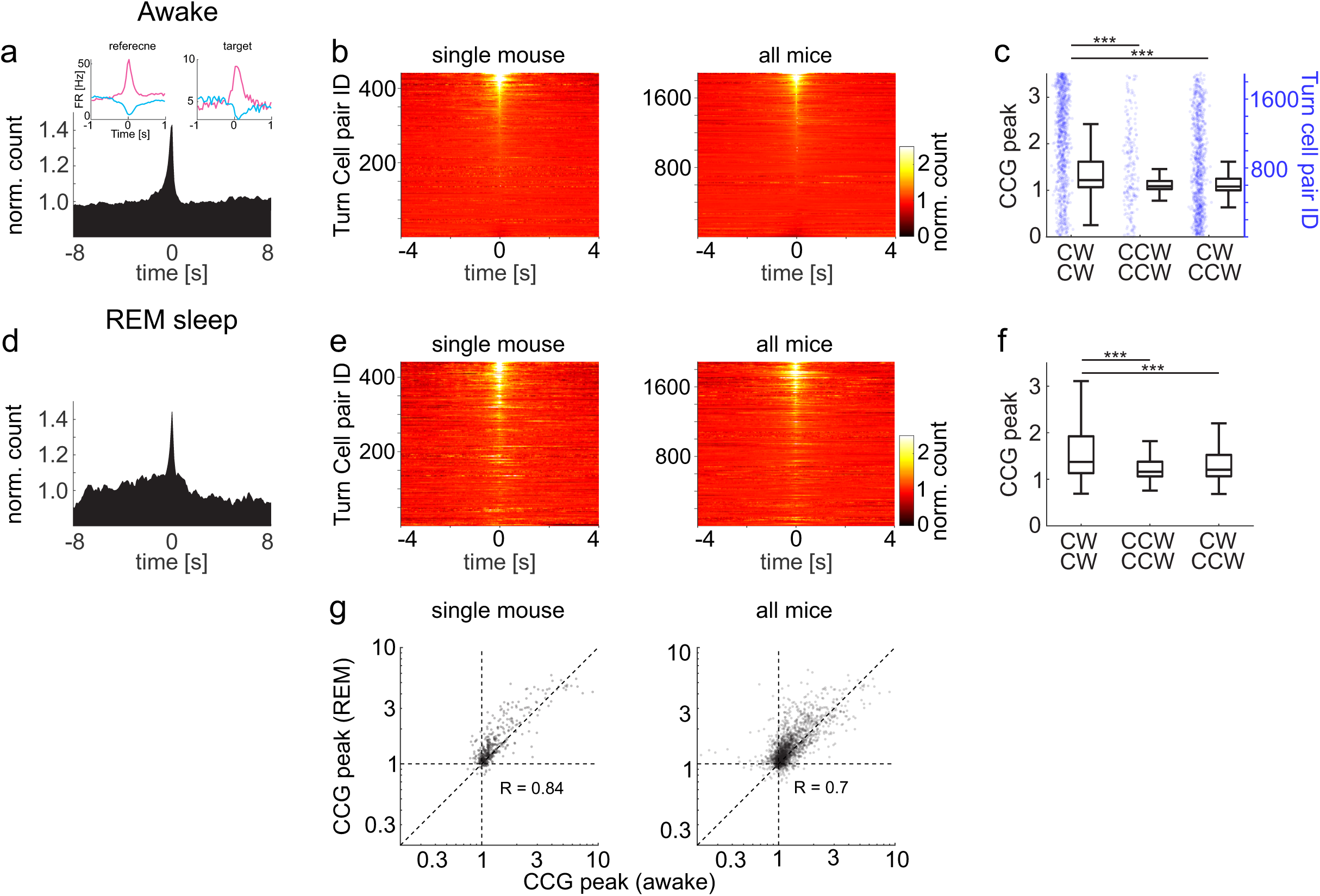
Correlational Structure of Turn Cells in Wakefulness and REM sleep. (a) Cross correlogram (CCG) of two example CW turn cells during wakefulness. The PTTH of the two cells (target and reference) are shown as insets for both CW (pink) and CCW (blue) turns. (b) CCG for turn cells pairs from an individual mouse (left; n = 441 pairs) or from all mice (right; n = 1888 pairs; 8 mice). The turn cell pairs are ranked based on the normalized value (norm. count) of the CCG at time 0. The pair with the smallest value is given ID 1. (c) Black: The CCG peak is the average normalized count of the CCG around time zero (± 30 ms). The box plots represent median, 1^st^ and 3^rd^ quartile and range of CCG peaks for CW-CW, CCW-CCW and CW-CCW pairs. Note that CW-CW pairs have a significantly larger CCG peak (p < 10^−10^ with Kruskal-Wallis test and p < 10^−19^ with Bonferroni post hoc test). Blue: The plot indicates, for each cell pair ID, the turn preferences of the cells within that pair (i.e. whether CW-CW, CCW-CCW or CW-CCW). Cell pair IDs are the same as in (B) right. Note that pairs with higher IDs are more likely composed of cells with CW-CW turn preferences while pairs with lower IDs are more likely made of cells with CW-CCW preferences. (d) CCG of the two example turn cells in (a) but during REM sleep. (e) As in (b) but during REM sleep. The cell pairs are ranked based on the ID from (b). (f) as in (c) but during REM sleep. Note that, as during wakefulness, CW-CW pairs have a significantly larger CCG peak o (p < 10^−10^ with Kruskal-Wallis test and p < 10^−19^ with Bonferroni post hoc test). (g) Scatter plot of CCG peaks during wakefulness plotted against REM sleep for a single example mouse (left; n = 441 pairs) and for all mice (right; n = 1888 pairs). Note the significant correlation in both plots (p < 10^−119^ for each).

**Extended Data Figure 3.**
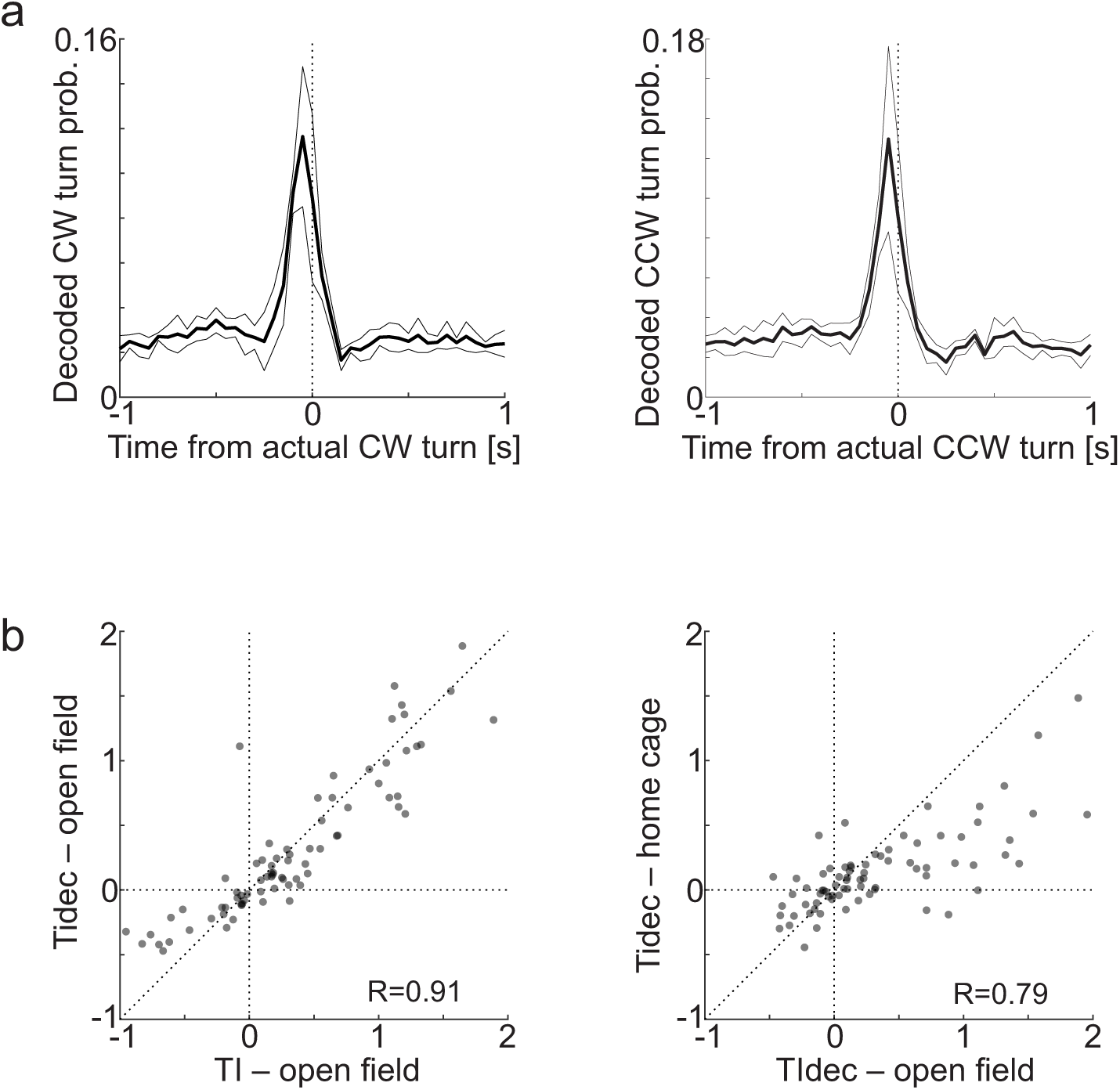
Decoded Turns and Decoded Turn Index. (a) Probability of detecting a turn from the decoded head direction (i.e. a decoded turn) around the time of an actual turn for CW (left) and CCW (right) turns (average of 1139±454 CW turns/mouse; average of 1080±404 CCW turns/mouse; n = 5 mice). . Thick line is average; Thin line is SD. The probability of decoding head turns within +/−200 ms from the onset of the actual head turn was 51.0% for CW and 53.5% for CCW turns. (b) Left: Scatter plot of Turn Indices of turn cells in SC computed from actual head turns (TI) against those computed from decoded head turns (TIdec) for periods during which the mice were in the open field arena. Note the tight correlation (p=1.3*10^−31^). Right: Same turn cells as in the left panel. Scatter plot of TIdec obtained during periods in the open field arena against TIdec obtained for periods in the home cage. Note that good correlation (P=5.4*10^−18^) despite the overall decrease in absolute TIdec values obtained in the home cage (78 cells; 5 mice).

**Extended Data Figure 4.**
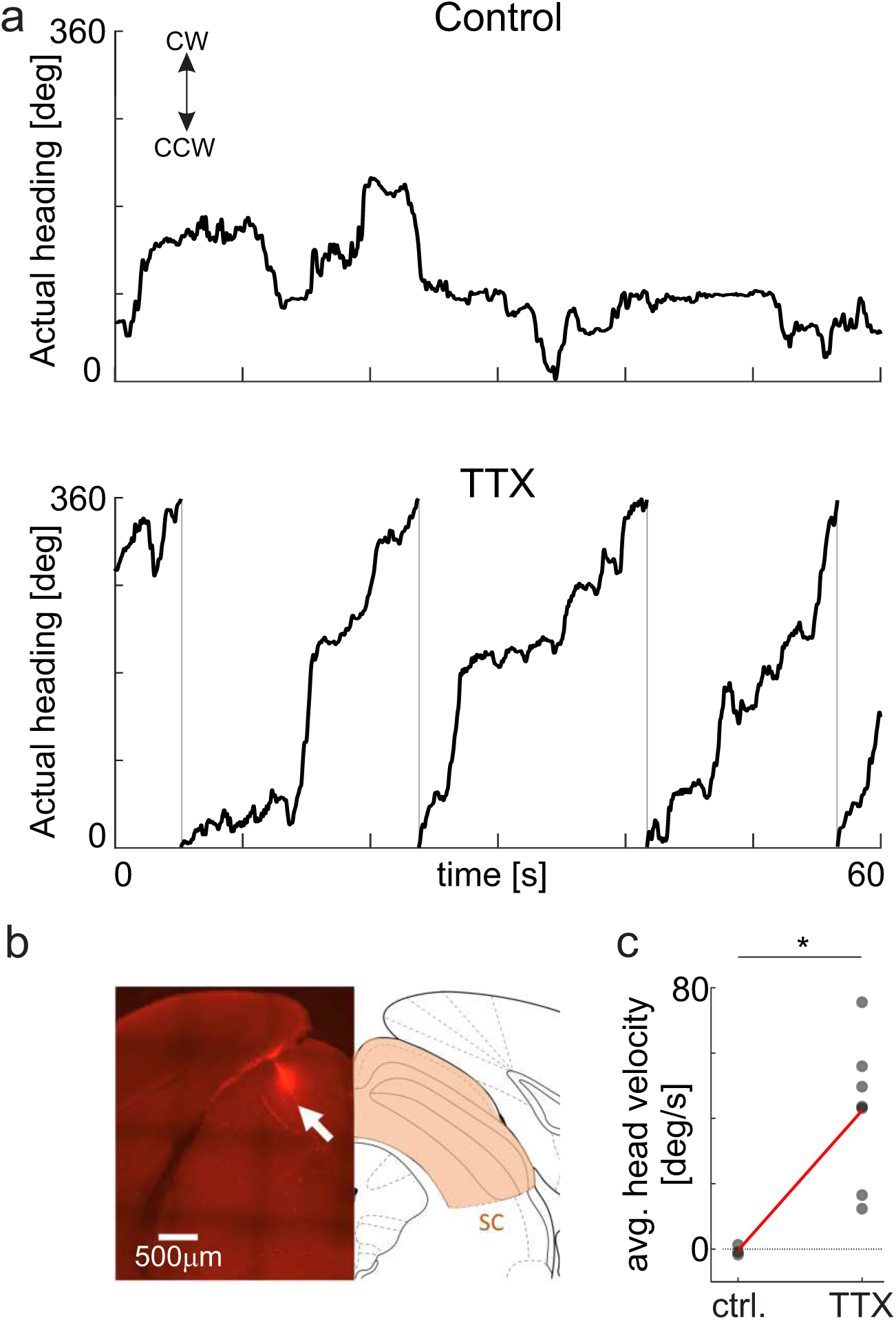
Impact of TTX in SC on actual heading. (a) Actual heading of an example untethered, awake animal under control conditions (top) and after injection of tetrodotoxin (TTX) in the right Superior Colliculus (SC; bottom). Upward changes in heading (i.e. positive slopes of the heading trajectories) are CW turns. Note that after injection of TTX the animals perform mainly CW turns (∼3.5 turns over a period of 60 s in this example). (b) Right panel: Photomicrograph of a coronal section of mouse brain illustrating the TTX injection site in the right SC (the TTX solution contained the red fluorescent dye Alexa 594). Left panel: Delineation of anatomical structures of corresponding coronal plane (Paxinos and Franklin, 2001. Bregma −3.88 mm). (c) Population data of averaged actual head velocity in control (ctrl.) versus TTX. Black data points: individual mice; Red: Population mean. Note the increase in positive velocities (i.e. CW turns) with TTX (p = 0.017; rank sum test). Control: 3 mice; TTX: 7 mice, including 2 control mice.

**Table S1.** The table provides experimental details relative to the animals and the recordings used throughout the study.

| mouse ID | 99 | 100 | 102 | 103 | 106 | 107 | 110 | 111 |
| --- | --- | --- | --- | --- | --- | --- | --- | --- |
| sex | M | M | M | M | M | M | M | M |
| Age @ implant [month] | 2.7 | 2.7 | 3.5 | 5.3 | 5.9 | 2.1 | 5.6 | 5.7 |
| Age @ recording [month] | 2.9 | 3 | 3.7 | 5.5 | 6.1 | 2.3 | 5.9 | 5.9 |
| Type of probe in SC | 1 shank (64-4) | 1 shank (64-4) | 1 shank (64-4) | 1 shank (64-4) | 1 shank (64-4) | 1 shank (64-4) | 1 shank (64-4) | 1 shank (64-4) |
| SC probe insertion angle toward medial [deg] | 24 | 24 | 24 | 24 | 20 | 20 | 20 | 20 |
| SC probe coordinate [mm, from Lambda] -- shank 1 | 0.0AP, -1.8LM | 0.0AP, -1.8LM | 0.0AP, -1.8LM | 0.0AP, -1.6LM | 0.0AP, -1.8LM | 0.0AP, -1.8LM | 0.0AP, -1.8LM | 0.0AP, -2.0LM |
| SC probe depth from brain surface [mm] | 1.22 to 2.48 | 0.65 to 1.91 | 0.77 to 2.03 | 0.44 to 1.70 | 1.07 to 2.33 | 0.86 to 2.12 | 1.44 to 2.70 | 1.99 to 2.25 |
| Type of probe in the Anterior Thalamus (AT) | 2 shank (64-3) | 2 shank (64-3) | 2 shank (64-3) | 2 shank (64-3) | 2 shank (64-3) | 2 shank (64-3) | 4 shank (64-1) | 2 shank (64-3) |
| AT probe insertion angle toward medial [deg] | 15 | 15 | 15 | 15 | 15 | 15 | 15 | 15 |
| AT probe coordinate [mm, from Bregma] -- shank 1 | -0.7AP, 1.3LM | -0.7AP, 1.3LM | -0.7AP, 1.3LM | -0.7AP, 1.5LM | -0.7AP, 1.5LM | -0.7AP, 1.5LM | -0.3AP, 1.5LM | -0.6AP, 1.5LM |
| shank 2 | -0.7AP, 1.55LM | -0.7AP, 1.55LM | -0.7AP, 1.55LM | -0.7AP, 1.75LM | -0.7AP, 1.75LM | -0.7AP, 1.75LM | -0.55AP, 1.5LM | -0.85AP, 1.5LM |
| shank 3 | --- | --- | --- | --- | --- | --- | -0.8AP, 1.5LM | --- |
| shank 4 | --- | --- | --- | --- | --- | --- | -1.05AP, 1.5LM | --- |
| AT probe depth from brain surface [mm] | 2.34 to 3.12 | 3.10 to 3.88 | 2.95 to 3.73 | 2.47to 3.25 | 2.47to 3.25 | 2.72 to 3.50 | 2.58 to 2.88 | 2.62 to 3.4 |
| Total number of units recorded in SC | 36 | 54 | 49 | 26 | 37 | 25 | 31 | 92 |
| Number of Turn Cells | 17 | 30 | 25 | 7 | 31 | 17 | 7 | 36 |
| Number of CW Turn Cells | 9 | 17 | 14 | 3 | 23 | 15 | 3 | 27 |
| Total number of units recorded in AT | 120 | 139 | 96 | 172 | 118 | 130 | 142 | 151 |
| Number of HD cells | 59 | 0 | 0 | 64 | 27 | 39 | 40 | 0 |
| HD decoding error [deg] | 9 | --- | --- | 10 | 12 | 9 | 9 | --- |
| Recording session [s] | 40920 | 41921 | 36527 | 36467 | 42964 | 80711 | 67359 | 72479 |
| Sleep duration [s] | 12740 | 14453 | 15586 | 17692 | 15674 | 41074 | 38334 | 40656 |
| REM sleep duration [s] | 944 | 643 | 1356 | 1729 | 1594 | 3662 | 3734 | 3070 |
| Number of REM sleep bouts | 16 | 6 | 22 | 33 | 28 | 48 | 49 | 40 |
| Number of head turns [CW / CCW] | 623 / 569 | 855 / 760 | 677 / 643 | 986 / 938 | 4290 / 4228 | 1842 / 1733 | 1175 / 1132 | 1932 / 1912 |

Datasets for Fig.4
| mouse ID | 83 | 85 | 112 | 115 | 116 | 117 | 118 | 119 |
| --- | --- | --- | --- | --- | --- | --- | --- | --- |
| sex | M | M | F | F | F | F | F | F |
| Age @ implant [month] | 3.4 | 3.7 | 3.4 | 4.9 | 12 | 12 | 12 | 12 |
| Age @ recording [month] | 3.9 | 4 | 5 | 5.8 | 12.2 | 12.5 | 12.7 | 12.9 |
| Type of probe in the Anterior Thalamus (AT) | 2 shank probe | 2 shank probe | --- | --- | 4 shank probe | --- | --- | 4 shank probe |
| AT probe insertion angle toward medial [deg] | 0 | 0 | --- | --- | 15 | --- | --- | 15 |
| AT probe coordinate [mm, from Bregma] -- shank 1 | -0.7AP, -0.7LM | -0.7AP, -0.7LM | --- | --- | -0.3AP, -1.4LM | --- | --- | -0.3AP, -1.4LM |
| shank 2 | -0.7AP, -0.95LM | -0.7AP, -0.95LM | --- | --- | -0.55AP, -1.4LM | --- | --- | -0.55AP, -1.4LM |
| shank 3 | --- | --- | --- | --- | -0.8AP, -1.4LM | --- | --- | -0.8AP, -1.4LM |
| shank 4 | --- | --- | --- | --- | -1.05AP, -1.4LM | --- | --- | -1.05AP, -1.4LM |
| AT probe depth from brain surface [mm] |  |  |  |  |  |  |  |  |
| Total number of units recorded in AT | 86 | 97 | --- | --- | 391 | --- | --- | 79 |
| Number of HD cells | 32 | 45 | --- | --- | 81 | --- | --- | 18 |
| HD decoding error [deg] | 8 | 8 | --- | --- | 10 | --- | --- | 9 |
| Recording session [s] (w/o TTX) | 30055 | 21945 | --- | --- | 18783 | --- | --- | 26257 |
| Sleep duration [s] | 12958 | 11309 | --- | --- | 6133 | --- | --- | 9417 |
| REM sleep duration [s] | 1140 | 865 | --- | --- | 213 | --- | --- | 229 |
| Number of REM sleep bouts | 20 | 16 | --- | --- | 3 | --- | --- | 4 |
| Recording session [s] (w/ TTX) | 21323 | 26401 | --- | --- | 41990 | --- | --- | 49428 |
| Sleep duration [s] | 14139 | 11138 | --- | --- | 35111 | --- | --- | 40562 |
| REM sleep duration [s] | 1454 | 758 | --- | --- | 1677 | --- | --- | 1119 |
| Number of REM sleep bouts | 13 | 8 | --- | --- | 23 | --- | --- | 18 |
| Time from open field exp. (no cable) to TTX inject. [hr] | --- | --- | 13.8 | --- | 5.9 | --- | --- | --- |
| average head velocity before TTX, no cable [deg/s] | --- | --- | -1.7 | -0.9 | 1.3 | --- | --- | --- |
| Time from TTX inject. to open field exp. (no cable) [hr] | 6.1 | 8 | 8 | --- | 11.7 | 14.6 | 13.4 | 13.7 |
| average head velocity after TTX, no cable [deg/s] | 43.1 | 43.6 | 16.5 | --- | 12.4 | 56 | 50 | 75.6 |

